# Molecular Insights into the Role of Water in Early-stage Human Amylin Aggregation

**DOI:** 10.1101/2022.08.29.505769

**Authors:** Ashley Z. Guo, Juan J. de Pablo

## Abstract

Human islet amyloid polypeptide (hIAPP or human amylin) is known to aggregate into amyloid fibrils and is implicated in the development of type II diabetes. Prefibrillar species in particular have been linked to cell loss, prompting detailed investigation of early-stage hIAPP aggregation. Insights into the mechanisms underlying early-stage aggregation and the key intermediate structures formed during aggregation are valuable in understanding disease onset at the molecular level and guiding design of effective therapeutic strategies. Here, we use atomistic molecular dynamics simulations with the finite temperature string method to identify and compare multiple pathways for hIAPP trimer formation in water. We focus on the comparison between trimerization from three disordered hIAPP chains (which we call “3-chain assembly”) and trimerization from an hIAPP dimer approached by a single disordered chain (called “2+1 assembly”). We show that trimerization is a process uphill in free energy, regardless of the trimerization mechanism, and that a high free energy barrier of 40 *k_B_T* must be crossed in 2+1 assembly compared to a moderate barrier of 12 *k_B_T* for 3-chain assembly. We find this discrepancy to originate from differences in molecular-level water interactions involved in the two trimerization scenarios. Furthermore, we find that the more thermodynamically favorable 3-chain assembly begins from a previously identified dimer intermediate exhibiting transient *β*-sheet character, which is then incorporated into a similar trimer intermediate, suggesting stepwise aggregation dynamics.

## I. INTRODUCTION

Abnormal aggregation of amyloidogenic proteins is implicated in numerous human diseases, including type II diabetes and various neurodegenerative diseases, such as Alzheimer’s disease. In each of these diseases, a particular protein self-assembles into a type of heavily *β*-sheet fibrillar aggregate known as amyloid.^1^ Human islet amyloid polypeptide (hIAPP or human amylin) is one such amyloidogenic protein; this 37-residue hormone is secreted with insulin in the pancreas and is involved in blood glucose regulation.^2,3^. Formation of amyloid aggregates of hIAPP has been linked to the development of type II diabetes as well as the loss of pancreatic *β*-cells,^4^ which has motivated the study of hIAPP aggregates and the mechanism through which they are formed.

The structure of the mature hIAPP fibril has been studied extensively via various structural characterization methods, including solid-state NMR (ssNMR) experiments, used to identify the how individual hIAPP monomers are arranged within a mature fibril. In this ssNMR model, hIAPP monomers are stacked one by one in the direction of the fibril axis, with each individual hIAPP chain in a U-shaped conformation with a region of *β*-strand on either side (in residues 8–17 and 28–37). As the hIAPP monomers stack in along the fibril axis, they form parallel *β*-sheets as adjacent U-shaped monomers align alongside each other. Two-dimensional infrared spectroscopy (2D-IR) experiments support this proposed structure,^5^ as do electron paramagnetic resonance (EPR) experiments.^6^ Additional experimental and computational studies have investigated the behavior of amylin and its mutants in the presence of various inhibitors and metals,^7–10^ as well as interactions with other amyloid-forming proteins or with membranes.^8,11–17^ Furthermore, studies of amylin have extended to examining structural rearrangements during aggregation,^18^ as well as identifying aggregation mechanisms.^19^

While mature amylin fibrils have been found to be biologically inert, previous studies have found early-stage aggregates to be associated with cytotoxicity.^20–23^ Prefibrillar species such as dimers, trimers, or higher order oligomers have been proposed as the key species responsible for inducing damage to cell membranes and eventually triggering cell death.^5,21,24–26^ Additionally, experiments in which cells undergo addition of hIAPP reveal membrane leakage prior to the formation of mature fibrils^27^ and disruption of cell membranes in regions separate from areas of fibril growth,^28^ further supporting the link between prefibrillar species and cellular damage.

A thorough investigation of early-stage fibril formation, including the mechanisms underlying amylin aggregation, is therefore critical in building a better understanding of how hIAPP behaves during the onset of disease and whether intermediate structures involved in the aggregation process could potentially be targeted therapeutically. hIAPP residues 20–29 have specifically been proposed to be key in amyloid formation,^29^ and mutations to this sequence lead to suppressed amylin aggregation.^29–32^ While residues 20–29 are not located in the parallel *β*-sheet regions formed in the mature hIAPP fibril, transient parallel *β*-sheets have been shown via 2D-IR experiments and molecular dynamics (MD) simulations to form in this region prior to fibril formation, particularly in residues 23–27 (with sequence `FGAIL`).^1,33^ Additionally, 2D-IR experiments with dihedral indexing have identified residues L12A13 to form a transient stacked turn or disordered *β*-sheet during aggregation.^34^

Multiple studies have utilized MD simulations to probe the dimerization of two hIAPP monomers into a U-shaped dimer,^17,33,35^ including our recent work^19^ employing the string method to discover the dimerization mechanism and confirming the formation of transient *β*-sheet in residues 12–13 and 20–29. The string method-based study found the final U-shaped dimer to be less thermodynamically stable than the disordered dimer by approximately 4.5 *k_B_T*, with a single major free energy barrier of 7 *k_B_T* in the dimerization process associated with formation of an intermediate structure exhibiting transient *β*-sheet in the aforementioned residues 12–13 and 20–29.^19^

Although these new insights have clarified the dimerization process, many key questions remain with regards to how higher order aggregates are formed, as well as how the unfavorable dimerization of amylin monomers fits into the amylin aggregation process at large. These questions include whether further addition of hIAPP monomers to the growing fibril proceeds similarly to the dimerization mechanism, and whether that process continues to be uphill in free energy. The recent application of the string method to the study of early-stage amylin aggregation has paved the way for studying these higher order aggregates; previous MD-based work on early-stage aggregation have largely been limited to study of the dimer, due to the reliance on the more computationally costly metadynamics approach. In this work, we tackle the next frontier in the early-stage aggregation process–trimerization–by extending the string method approach to discover multiple possible trimerization mechanisms and compare their respective thermodynamic properties. We focus specifically on the comparison of trimerization from three disordered amylin chains versus trimerization from a disordered chain approaching an amylin dimer, the distinctly different free energy barriers encountered in each case, and most interestingly, the role of molecular interactions with water that underlie the key differences between the two aggregation processes.

## II. RESULTS AND DISCUSSION

The finite temperature string method was used, as detailed in the Methods section, to identify and investigate aggregation mechanisms for hIAPP trimerization. We perform the finite temperature string method using two collective variables: (1) a measure of parallel *β*-sheet character, 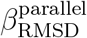, which is described in the Methods, and (2) the radius of gyration (*R_g_*) of the three protein chains. 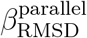 provides a continuous measure of the amount of parallel *β*-sheet formed, while *R_g_* provides a measure of compactness; we use them together to characterize the trimerization process from a disordered state with little *β*-sheet to a more compact aggregated state exhibiting high *β*-sheet secondary structure.

### A. Trimer Assembly from the Disordered State (“3-chain Assembly”)

We begin by investigating assembly of the hIAPP trimer from three separate disordered hIAPP chains; we refer to this assembly process as “3-chain Assembly”. Figure 1 shows the trimerization pathway discovered via the finite temperature string method. One end of the pathway corresponds to the disordered state of the hIAPP trimer (*R_g_* = 2.16, 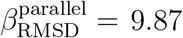, shown in Panel I), while the opposite end corresponds to the fully formed trimer (*R_g_* = 2.99, 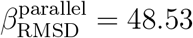, shown in Panel IV). Representative snapshots highlighting key conformational changes along the trimerization pathway are shown at the top of Figure 1.

**FIG. 1.**
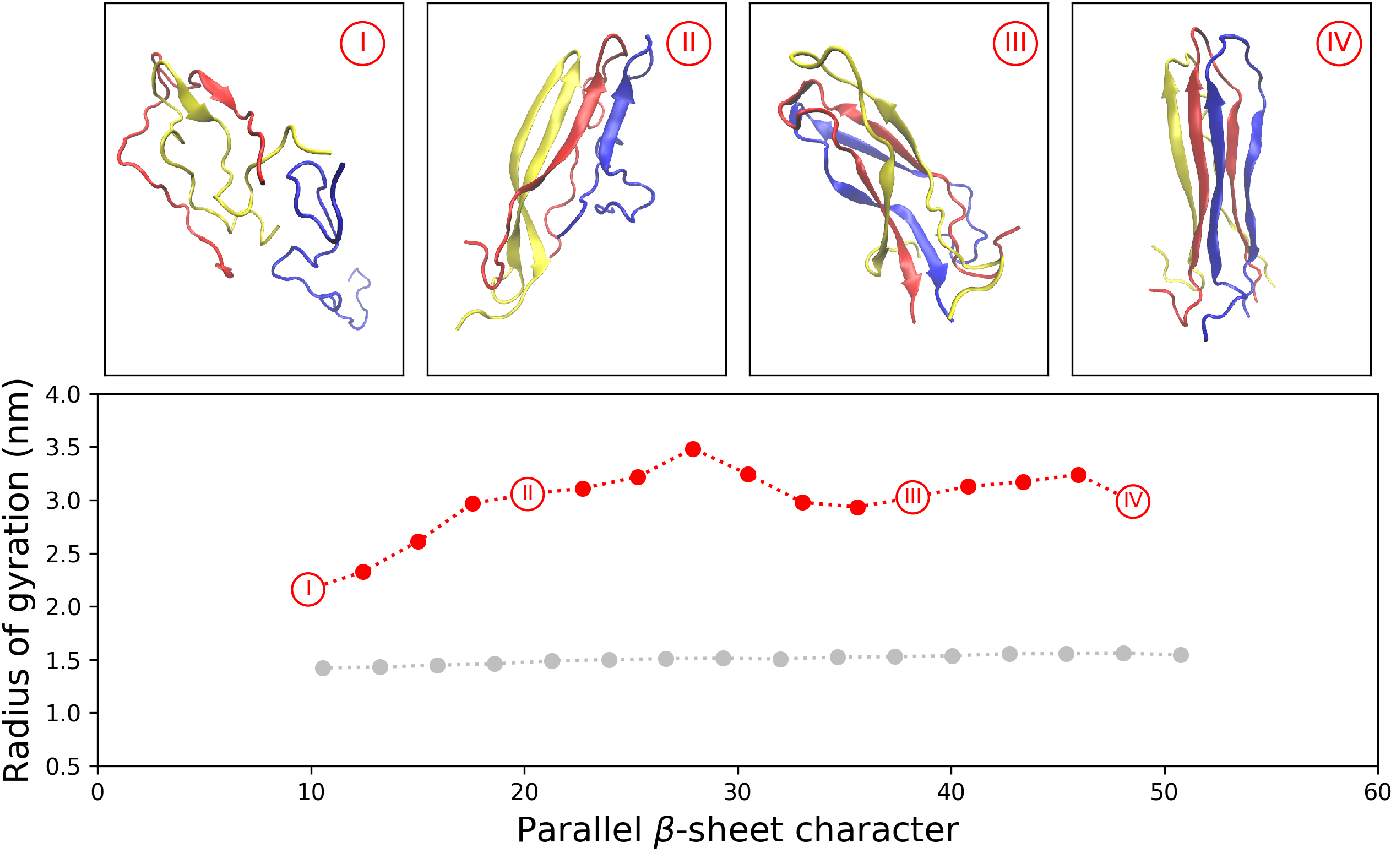
3-chain trimerization pathway calculated from the finite temperature string method, with initial configurations input into the string method shown in grey. Four representative snapshots show mechanistic details during trimer formation. Water and counterions are removed for clarity. The disordered end of the string (Panel I) includes the previously studied dimer intermediate (formed by the yellow and red chains). A similar intermediate comprised of all three chains is discovered and shown in Panel II.

In Panel I, a disordered amylin chain exhibiting no secondary structure approaches two amylin chains that are loosely associated. Interestingly, the structure of these two chains corresponds to the intermediate structure previously observed for hIAPP dimerization,^19^ with parallel *β*-sheet formed in the turn region suspected to play a key role in amyloid aggregation.^29^ Panel II shows that the disordered amylin chain has associated with the dimer intermediate observed in Panel I, forming a similar trimer intermediate that also exhibits parallel *β*-sheet, this time in the bend region across all three chains. Taken together, Panels I and II suggest that formation of higher-order hIAPP aggregates in the 3-chain assembly scenario proceeds in a stepwise manner, with individual amylin chains added sequentially to a growing aggregate at the intermediate parallel *β*-sheet stage. Increasing amounts of parallel *β*-sheet are formed between the three amylin chains through Panels III and IV, leading to the formation of the full trimer, shown in Panel V.

Free energy changes along this 3-chain assembly pathway were calculated as described in the Methods section and are shown in Figure 2. A single major free energy barrier of approximately 12 *k_B_T* is observed, corresponding to the formation of the intermediate *β*-sheet structure as indicated by the snapshots in Figure 2 and in Panel II of Figure 1. The amylin dimer intermediate rearranges to allow incorporation of the approaching disordered third amylin chain during this 12 *k_B_T* increase in free energy, followed by a slight drop in free energy after the trimer intermediate is formed.

**FIG. 2.**
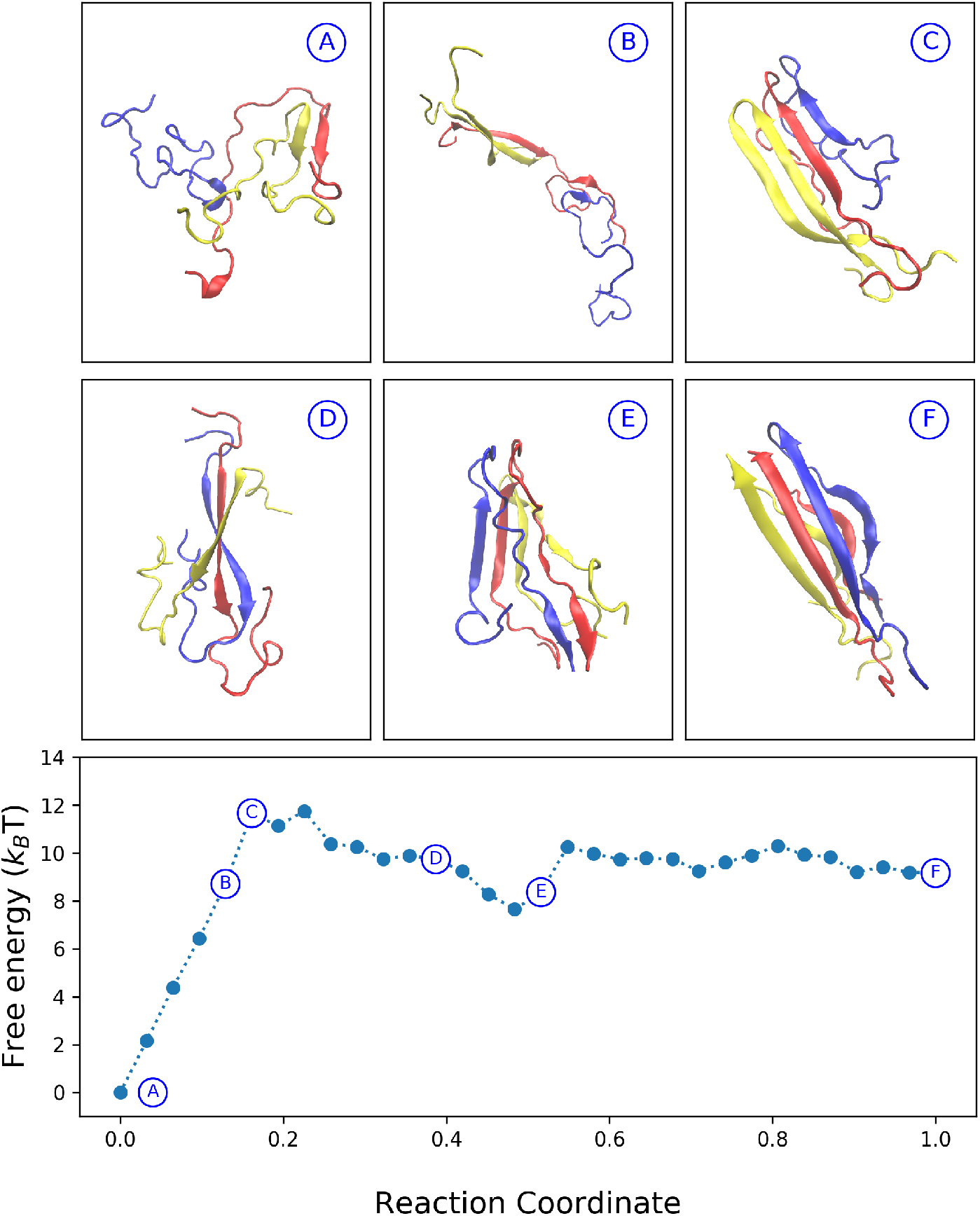
Free energy profile along the 3-chain trimerization pathway found via the finite temperature string method. The reaction coordinate extends from 0.0 (disordered state) to 1.0 (fully formed trimer). Free energy is calculated using the procedure described in the Methods, sampling each of the 32 cells for 150 ns each. Representative snapshots are shown to illustrate conformational changes during trimerization. A free energy barrier of approximately 12 *k_β_T* is found, corresponding to the formation of a transient *β*-sheet structure (shown in Panel C).

Figure 3 shows the changes in average secondary structure per residue over the 3-chain assembly process, calculated using the DSSP algorithm.^36^ Through comparison with Figure 2, it becomes clear that *β*-sheet is formed in the intermediate structures in residues 12–13 and 20–29, which have been previously proposed^29,34^ as regions exhibiting transient *β*-sheet during aggregation and observed to do so in studies of the hIAPP dimer.^19^ Figure 3 also shows *β*-sheet forming primarily in the C-termini before advancing to the N-termini, a pattern which was also observed in previous dimer studies.^19^

**FIG. 3.**
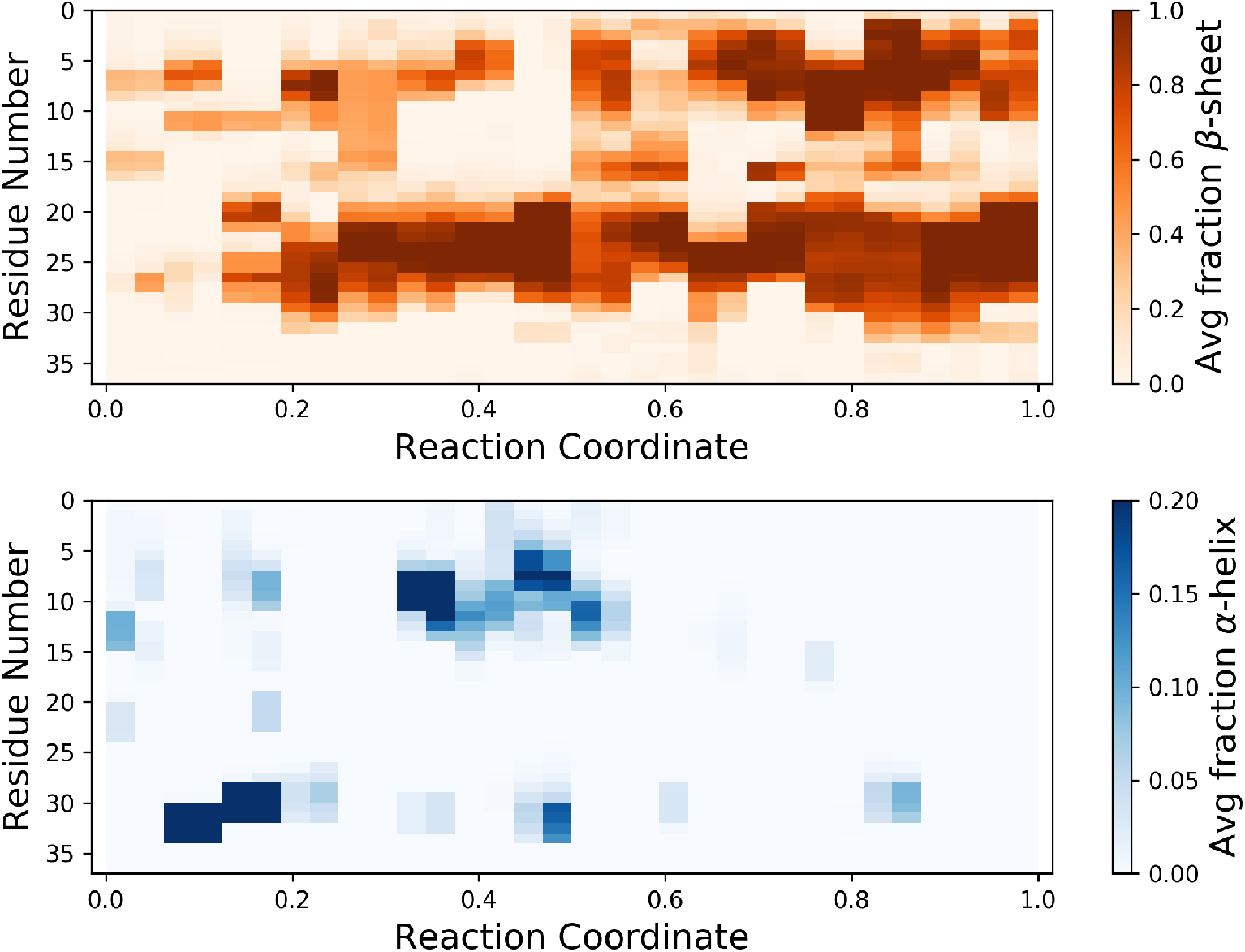
Average *α*-helix and *β*-sheet secondary structure along the 3-chain trimerization pathway, plotted by residue (ranging from 1 to 37, averaged over all three hIAPP chains). Secondary structure was calculated using the DSSP algorithm^36^ and averaged over 150 ns of sampling for each of the 32 bins along the reaction coordinate. *β*-sheet is found to transiently form in residues 12–13 and 20–29 during formation of the intermediate *β*-sheet structure, and as trimerization progresses, *β*-sheet extends to both termini, with more heavily *β*-sheet character observed in the C-termini.

### B. Trimer Assembly from the Dimer State (“2+1 Assembly”)

In addition to studying assembly of the hIAPP trimer from three disordered chains, we applied the finite temperature string method to the study of a trimer assembled from a single disordered amylin chain approaching a fully formed amylin dimer. We refer to this process as “2+1 Assembly”. The trimerization pathway discovered using the string method is shown in Figure 4; the disordered end of the pathway is found at (*R_g_* = 2.62, 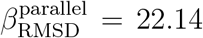, shown in Panel I), and the fully formed trimer is found at the opposite end of the discovered pathway at (*R_g_* = 2.98, 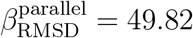, shown in Panel IV). Again, representative snapshots highlight the relevant conformational changes along the trimerization pathway, shown at the top of Figure 4. Panels I and II show that the hIAPP dimer is first stabilized, before the disordered third amylin chain is gradually incorporated in Panels III and IV. In contrast with 3-chain assembly and previous studies of the dimer,^19^ there is no clear intermediate structure exhibiting *β*-sheet in the bend region.

**FIG. 4.**
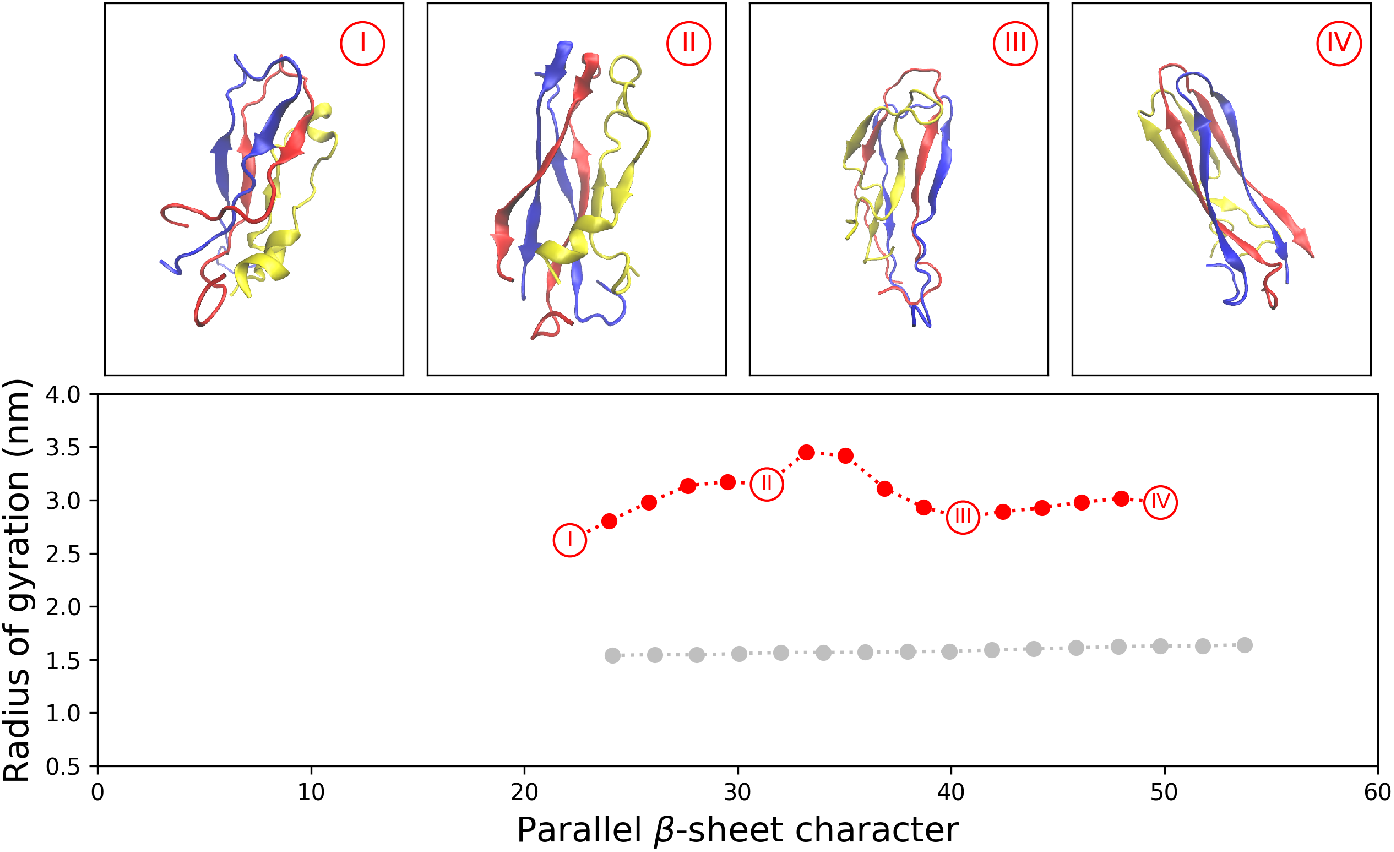
2+1 assembly pathway calculated from the finite temperature string method. Grey points show initial configurations input into the string method. Four representative snapshots illustrate conformational changes observed during trimer formation. Water and counterions are not shown. The disordered end of the string (Panel I) shows a loosely formed dimer and a disordered third chain. Increased *β*-sheet is formed gradually until the full trimer is formed (Panel IV).

We also calculate the free energy profile along the 2+1 transition pathway, shown in Figure 5. A steady increase of approximately 40 *k_B_T* is calculated for the 2+1 assembly process, with no clear intermediate metastable state. This 40 *k_B_T* rise in free energy is associated with the stabilization of the hIAPP dimer and initial addition of the third hIAPP chain to the dimer, as shown in Panels A-D. The free energy fluctuates around a steady value as the third chain gradually forms greater amounts of *β*-sheet with the dimer, eventually forming the full hIAPP trimer, shown in Panels E and F.

**FIG. 5.**
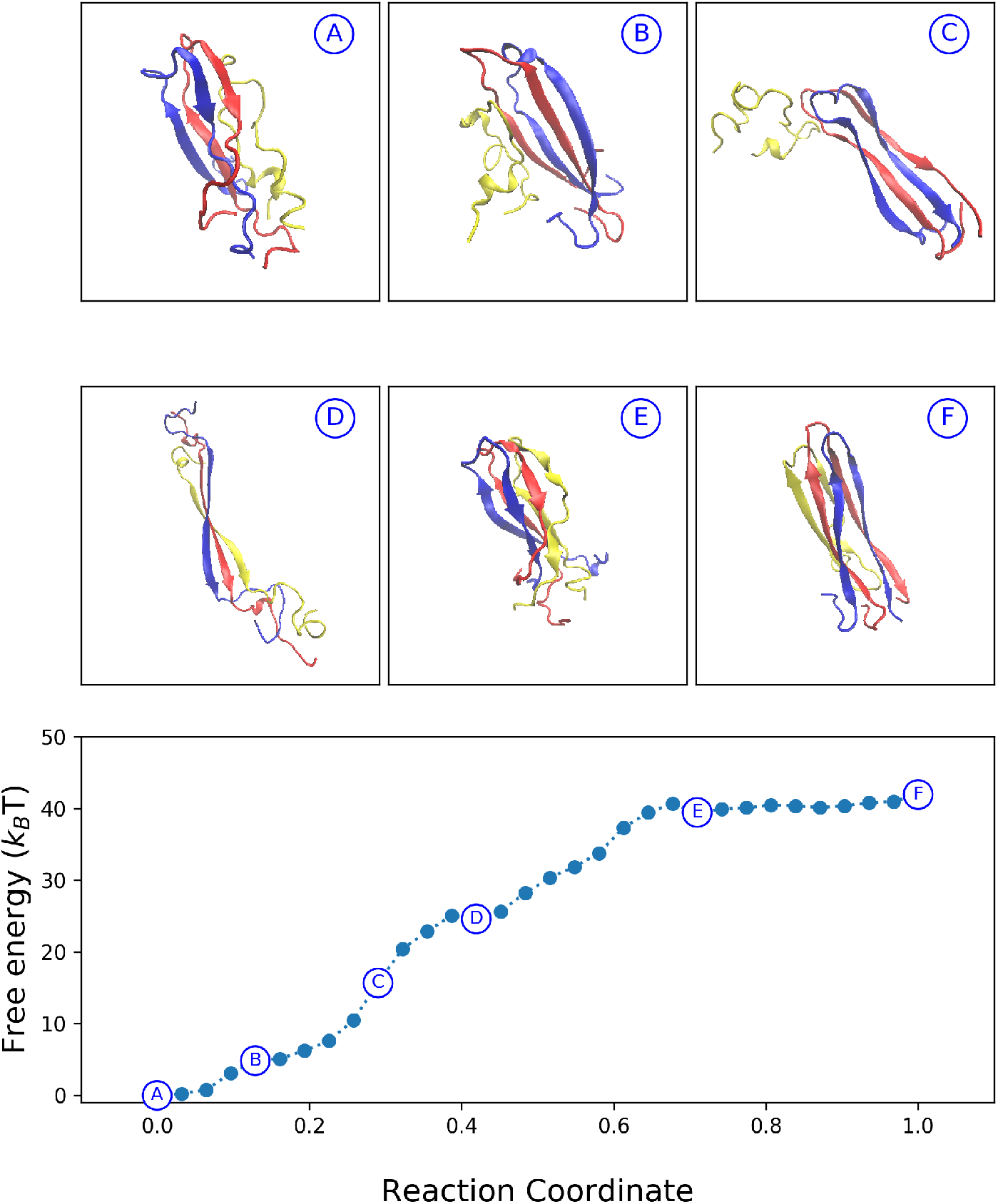
Free energy profile along the 2+1 assembly pathway found via the finite temperature string method. The reaction coordinate proceeds from 0.0 (disordered state containing a loosely formed dimer and third disordered chain) to 1.0 (fully formed trimer). Free energy is calculated using the procedure described in the Methods, sampling each of the 32 cells for 150 ns each. Representative snapshots are shown to illustrate conformational changes during trimerization. A free energy barrier of approximately 40 *k_β_T* is found. The intermediate transient *β*-sheet structure observed during 3-chain assembly is not found.

Following our analysis for 3-chain assembly, we now calculate the average secondary structure per residue for the 2+1 assembly process with the DSSP algorithm.^36^ Figure 6 shows these results, with unsurprising outcomes: compared to 3-chain assembly, 2+1 assembly begins with greater amounts both *β*-sheet (from the already-assembled dimer) and *α*-helix (from the approaching third chain). *β*-sheet gradually increases across the course of 2+1 assembly, with *β*-sheet forming earlier and more heavily in the C-termini before the N-termini, which was also observed for 3-chain assembly.

**FIG. 6.**
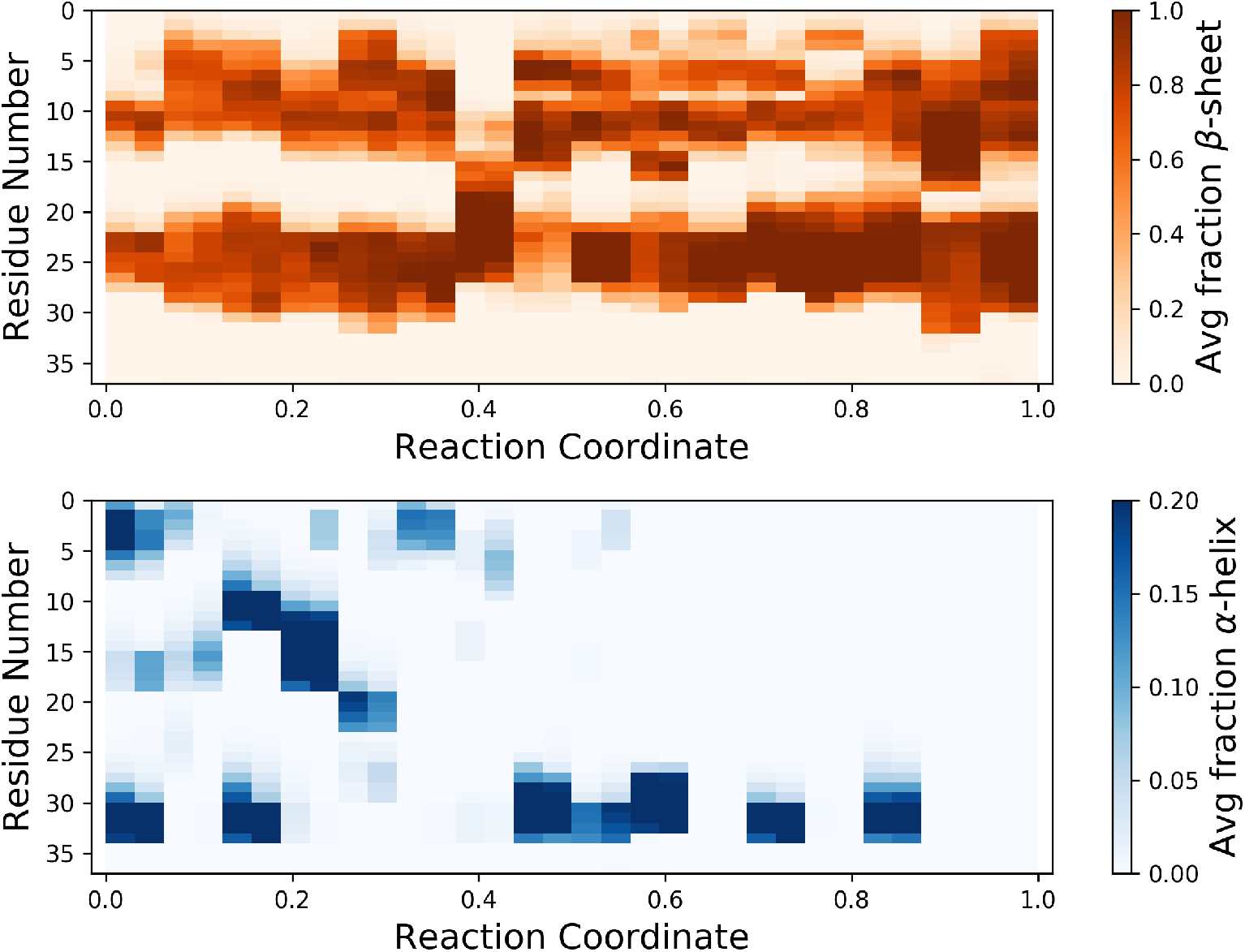
Average *α*-helix and *β*-sheet secondary structure along the 2+1 assembly pathway, plotted by residue (ranging from 1 to 37, averaged over all three hIAPP chains). Secondary structure was calculated using the DSSP algorithm^36^ and averaged over 150 ns of sampling for each of the 32 bins along the reaction coordinate. As with 3+1 assembly, *β*-sheet extends to both termini, with more heavily *β*-sheet character observed in the C-termini.

### C. “3-chain Assembly” versus “2+1 Assembly”

In order to understand the differences in mechanistic details and the discrepancy between the thermodynamics of the two assembly processes (12 *k_B_T* free energy barrier for 3-chain assembly vs 40 *k_B_T* for 2+1 assembly), we perform a series of comparisons to uncover the key differences between the two trimerization pathways: (1) calculation of protein-water and protein-protein hydrogen bonds during trimerization; (2) decomposition of free energy into entropic and enthalpic components; and (3) decomposition of potential energy contributions into inter- and intra-chain components. These three comparisons will allow us to isolate and contrast specific interactions that contribute to the previously calculated free energy profiles, thereby identifying how the two mechanisms differ and what contributes the unfavorability of 2+1 assembly compared to 3-chain assembly.

We begin by comparing protein-water and protein-protein H-bonds formed during the trimerization process, shown in Figure 7. Both assembly processes show a steady rise in protein-protein hydrogen bonds as the full trimer is formed. As expected, 2+1 assembly begins with higher protein-protein H-bond count than 3-chain assembly, reflecting the heavily *β*-sheet dimer required for the 2+1 pathway. Protein-water H-bonds fluctuate during both trimerization processes; however, there are three distinct downward trends in protein-water H-bonds observed during 3-chain assembly, which is not observed in 2+1 assembly. This is especially distinct during the formation of the full trimer in the last fifth of the 3-chain assembly process, indicating the loss of protein-water H-bonds while protein-protein H-bonds are gained during formation of the full trimer.

**FIG. 7.**
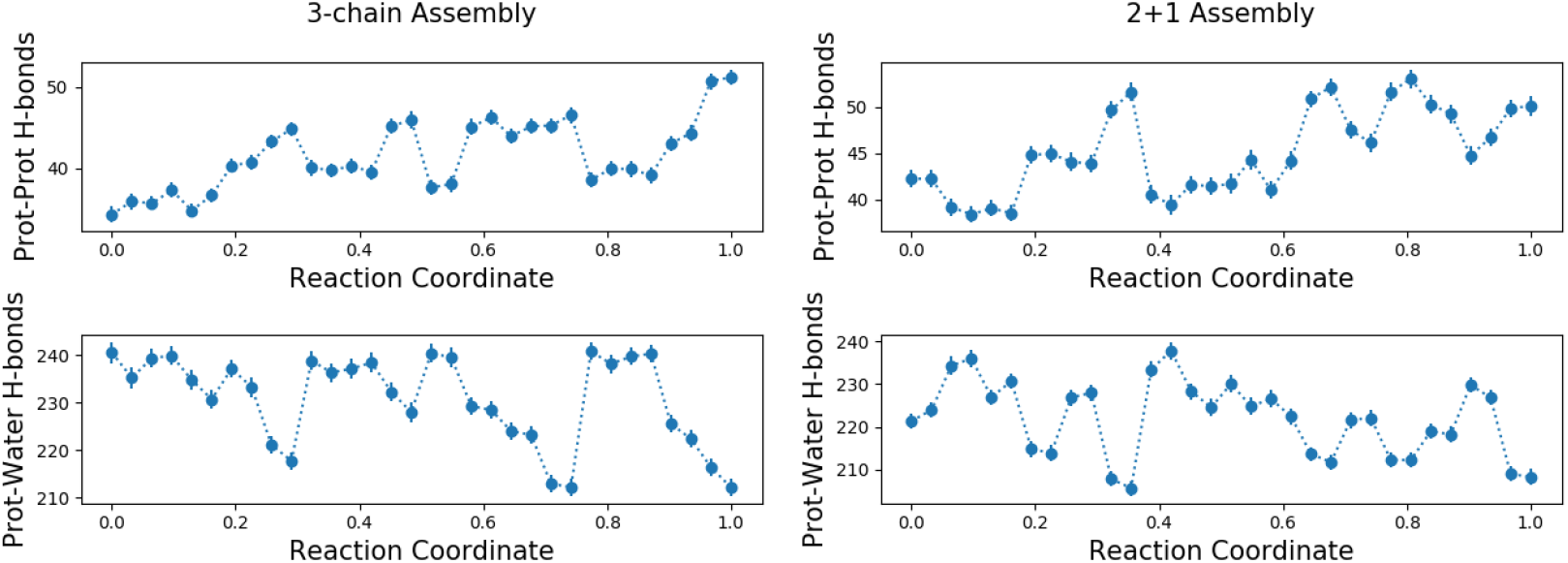
Average protein-protein and protein-water hydrogen bonds within 3 Åover the course of dimerization for both trimerization processes, shown with standard error. Average number of H-bonds are calculated from each of the 32 cells sampled during free energy calculation, from 15003 snapshots per cell, using the GROMACS `g_hbond` tool. While protein-protein H-bonds trend upward during both assembly processes, the protein-water H-bonds show a steeper decrease in 3-chain assembly compared to 2+1 assembly. Protein-protein H-bonds are found to form while protein-water H-bonds are lost during 3-chain assembly; this is not observed for 2+1 assembly.

The protein-water and protein-protein H-bond profiles shown in Figure 7 were then compared with profiles of enthalpic and entropic changes along the two trimerization pathways, shown in Figure 8. Free energy profiles for both assembly processes were split into entropic and enthalpic contributions. Entropic contributions were calculated using Δ*A* = Δ*U* – Δ*TS*, using potential energy differences Δ*U* and free energy differences Δ*A* for each cell sampled along each trimerization pathway, calculated with respect to the first disordered state bin.

**FIG. 8.**
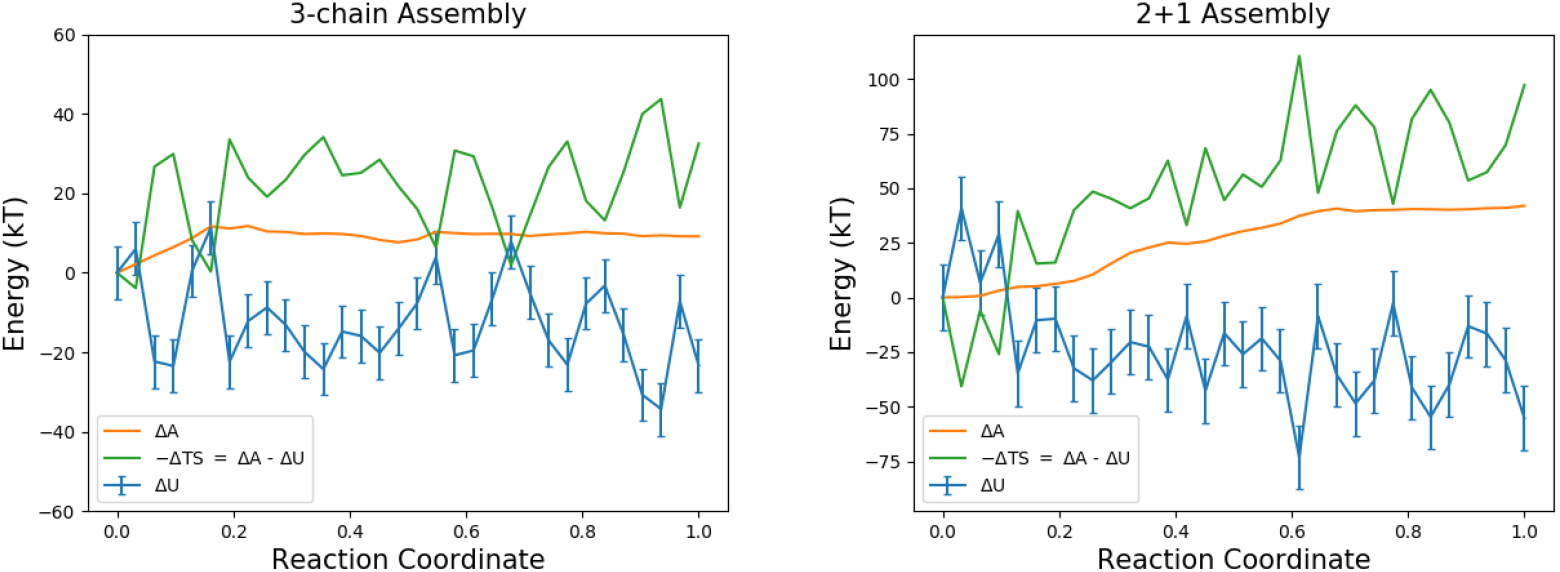
Decomposition of free energy into enthalpic (Δ*U*) and entropic (–Δ*TS*) contributions for both trimerization mechanisms. Δ*A* is taken from the free energy calculation, performed as described in Methods and shown in Figures 2 and 5. Average potential energy is calculated from each of the 32 cells sampled during free energy calculation, from a total of 15003 snapshots per cell. Entropic contributions are then calculated as – Δ*TS* = Δ*A* – Δ*U*.

Enthalpic and entropic contributions are both observed to fluctuate throughout 3-chain assembly, with minima in the enthalpic term corresponding with peaks in entropy across the trimerization process. While the free energy profile for 3-chain assembly does not indicate that the fully-formed trimer is metastable, a final dip in the enthalpic term along with its corresponding peak in the entropic term at the end of trimerization suggest there is some degree of entropic stabilization, which occurs in the same period in which protein-protein H-bonds are found to increase at the expense of protein-water H-bonds, shown in Figure 7.

In contrast, 2+1 assembly begins with an initial rise in potential energy and decrease in entropy. This feature is not observed in the free energy decomposition for 3-chain assembly, suggesting that the 2+1 assembly process must undergo a transition through enthalpically unfavorable conditions that are not observed during 3-chain assembly. As trimerization progresses after the third disordered chain meets the already-formed dimer structure, the entropic contribution steadily rises, leading to a steady and substantial rise in free energy of approximately 40 *k_β_T*, which was previously shown in Figure 5.

Additional insights into the differences between 3-chain assembly and 2+1 assembly were found by further decomposing the potential energy profiles for the two assembly processes into intrachain, interchain, and chain-water contributions. The stark differences in trends between the two assembly processes are shown in Figure 9. In 3-chain assembly, intrachain potential energy steadily rises throughout trimerization, while in 2+1 assembly it steadily decreases. Meanwhile, interchain potential energy fluctuates during 3-chain assembly before finally stabilizing during formation of the full trimer, in the same period of time where protein-water H-bonds sharply decreased while protein-protein H-bonds sharply increased (Figure 7). In contrast, interchain potential energy is initially stable for 2+1 assembly, rises during trimerization, and fluctuates during formation of the full trimer. Chain-water potential energy appears to fluctuate independently of inter- and intrachain interactions during 3-chain assembly, with a slight upward trend as the trimer forms; in 2+1 assembly, this chain-water potential energy exhibits a slight downward trend, with dips in chain-water interactions corresponding to peaks in intrachain interactions during the first half of trimerization, and with interchain interactions during the last half.

**FIG. 9.**
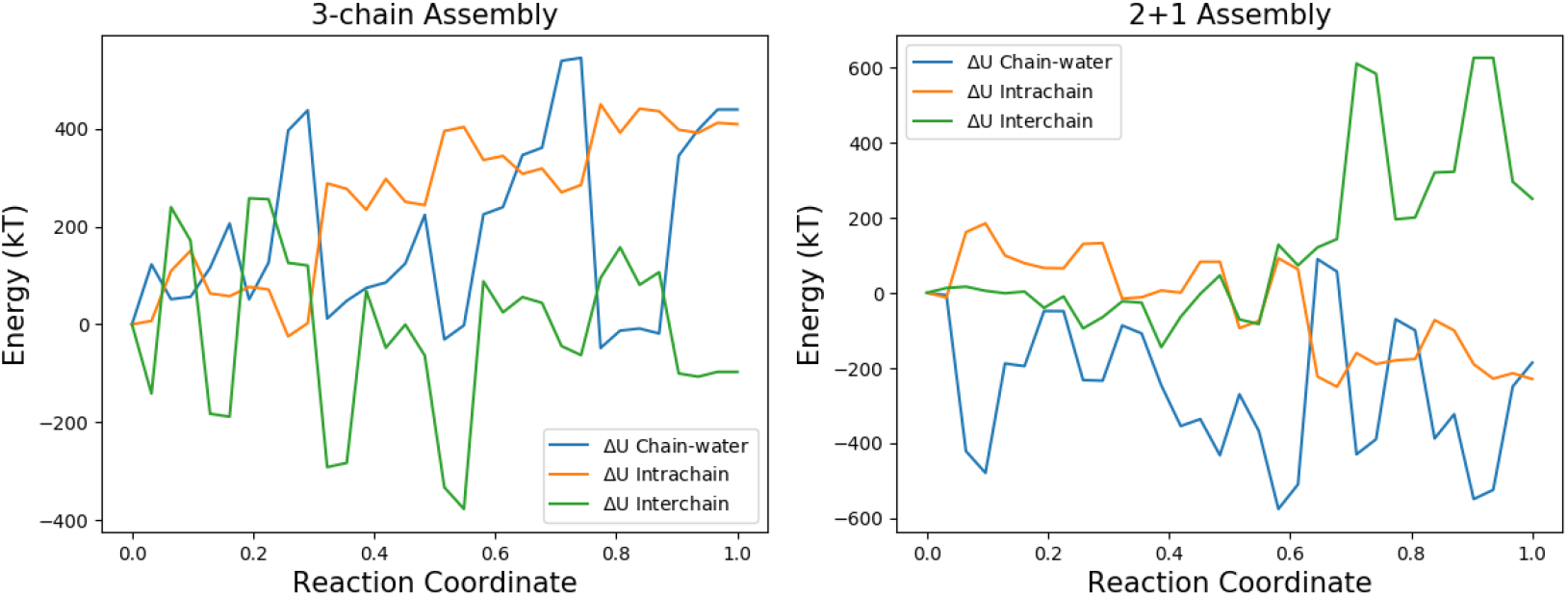
Decomposition of potential energy into interchain, intrachain, and chain-water interactions for both trimerization mechanisms. Energies are calculated from each of the 32 cells sampled during free energy calculation, from a total of 15003 snapshots per cell. Note the contrasting trends between the two assembly processes for all three interactions plotted.

Taken together, the comparisons above indicate that the role of water in each trimerization process contributes greatly to the distinct differences between 3-chain and 2+1 assembly; specifically, the pre-formed dimer in 2+1 assembly is stabilized by its protein-water interactions, and breaking of these protein-water interactions in order to form new protein-protein contacts during trimerization becomes unfavorable compared to formation of protein-protein contacts from three disordered chains. Decreases in protein-water H-bonds tend to correspond to gains in protein-protein H-bonds in 3-chain assembly, but this is not observed during 2+1 assembly. Increased enthalpic and decreased entropic contributions to free energy are observed in the first portion of 2+1 assembly, when the pre-formed dimer and disordered third chain are still separated and each surrounded by water; these conformations are not observed at any time during the 3-chain assembly process, and neither are these thermodynamic features. A sharp rise in entropic contributions and fall in enthalpic contributions, however, is observed at the very end of 3-chain assembly, corresponding to the final simultaneous drop in protein-water H-bonds and gain in protein-protein H-bonds; this, in turn, is not observed at any time during the 2+1 assembly process. Furthermore, chain-water potential energy decreases during 2+1 assembly, with minima corresponding to peaks in either intra- or interchain potential energy, indicating a tendency toward stabilization of protein-water interactions during 2+1 assembly at the expense of establishing energetic stabilization of protein-protein interactions within the forming amylin trimer. As the pre-formed dimer and disordered third chain in 2+1 assembly are brought together in water, the stabilization of pre-existing protein-water contacts is prioritized before the establishment and stabilization of new interactions between all three chains, reflected in the thermodynamic quantities discussed and compared here. We further confirm that it is the molecular interactions involving water that drive the differences observed in 3-chain and 2+1 assembly by recalculating the free energy profile of trimerization for both processes (as described in the Methods) with explicit water molecules replaced by the OBC GBSA implicit water model.^37^ With every individual interacting water atom now replaced with a dielectric continuum, the free energy profiles are considerably flattened, as displayed in Figure 10, suggesting that the key differences between 3-chain and 2+1 assembly indeed originate from molecular-level interactions with water.

**FIG. 10.**
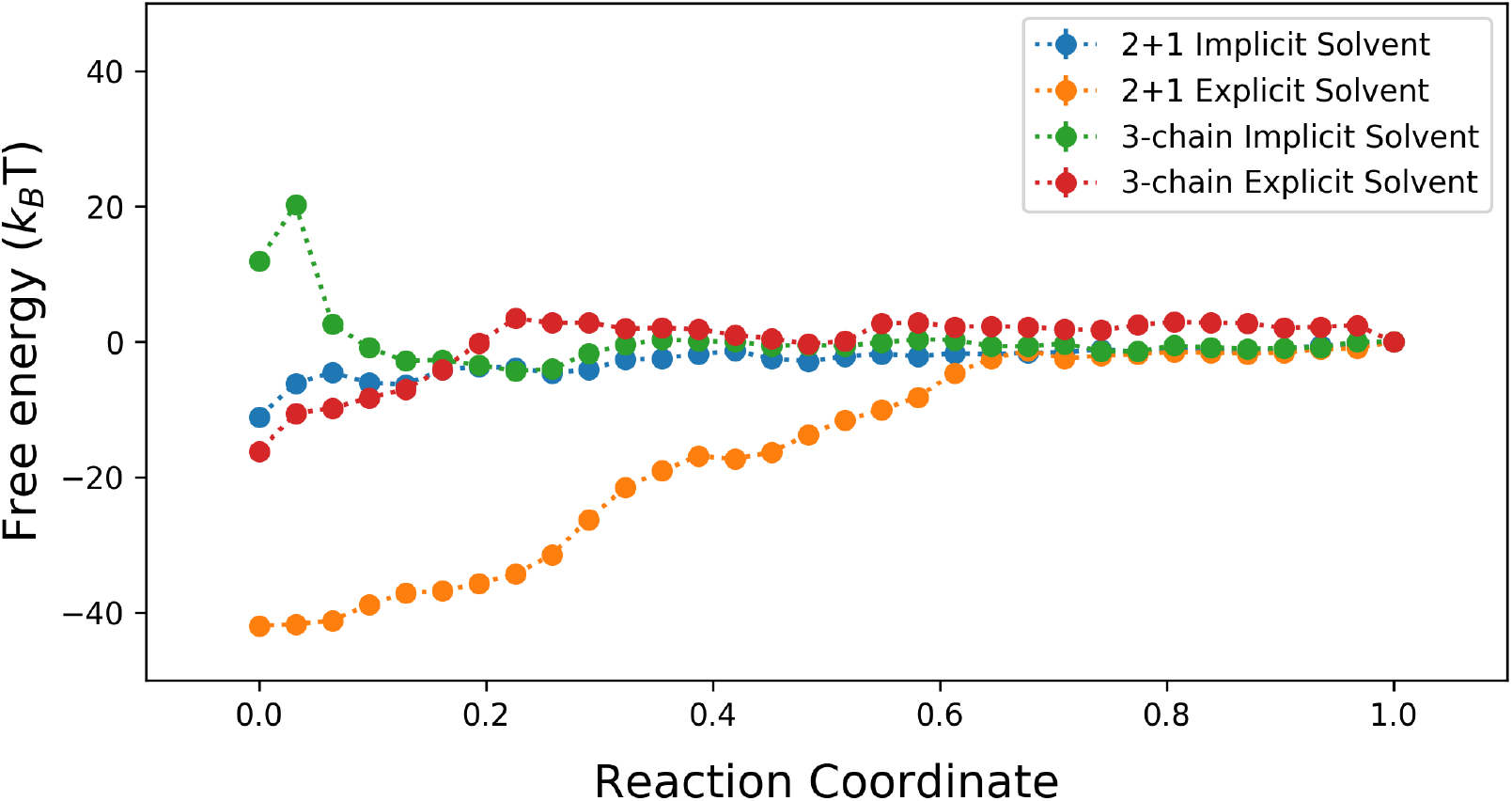
Comparison of free energy profiles for both 3-chain and 2+1 assembly, using an explicit water model (as was shown in Figures 2 and 5) versus implicit water. When the explicit water interactions are replaced with a dielectric continuum as in the implicit model, the free energy profiles are noticeably flattened, suggesting that the moleuclar-level interactions with water are responsible for the free energy barriers originally observed. Free energy profiles are aligned for all four profiles at the fully-formed trimer.

## III. CONCLUSIONS

A finite-temperature string method approach was used to study multiple pathways of hIAPP trimer formation and their thermodynamics, with specific focus on assembly from three disordered chains (“3-chain assembly”) versus assembly from a dimer and one disordered chain (“2+1 assembly”). In both 3-chain assembly and 2+1 assembly, the fully-formed trimer was found to lie in a global free energy minimum, separated from the fully-formed dimer by a climb in free energy. This climb is approximately 12 *k_B_T* for 3-chain assembly, and a steep 40 *k_B_T* for 2+1 assembly; neither fully-formed trimer structure is found to be metastable.

For 3-chain assembly, crossing of the 12 *k_B_T* barrier corresponds to formation of an intermediate structure exhibiting parallel *β*-sheet in residues 12–13 and 20–29 across all three hIAPP chains, a structure which has been previously proposed and similar to the intermediate *β*-sheet structure observed in computational studies of the hIAPP dimer. Interestingly, the string method identifies the disordered end of the 3-chain assembly as a conformation which includes this dimer intermediate state, suggesting that hIAPP aggregation via 3-chain assembly proceeds in a stepwise manner, using the intermediate *β*-sheet structure as a template for fibril propagation.

Furthermore, a series of comparisons was used to investigate the nature of the stark difference between the 12 *k_B_T* 3-chain assembly process and the 40 *k_B_T* 2+1 assembly process. Analysis of protein-water and protein-protein H-bonds over trimerization, decomposition of free energy into enthalpic and entropic contributions, and decomposition of potential energy into interchain, intrachain, and chain-water interactions linked the key differences between 3-chain and 2+1 assembly to their differences in molecular-level interactions with water. The relatively high number of pre-existing chain-water interactions in 2+1 assembly compared to 3-chain assembly underlie the individual differences in entropic, enthalpic, and hydrogen bond contributions, which together ultimately result in the two distinctly different trimerization free energy profiles.

Although the current work demonstrates that 3-chain assembly is more thermodynamically favorable versus 2+1 assembly, both processes are uphill in free energy, along with the dimerization process studied in our previous work. However, aggregates have been demonstrated to form experimentally through seeding and incubation procedures. Based on our finding that systems with an increased number of protein-water H-bonds undergo less favorable aggregation, we hypothesize that aggregates can be stabilized as a result of increased competition for hydrogen bonding with water from other molecules or due to a densely concentrated environment. A forthcoming publication will investigate this further by introducing additional species into the system, including salts and readily H-bond-forming molecules.

Additionally, questions still remain on whether further growth toward higher order oligomers will remain uphill in free energy, and whether spontaneous fibril growth only takes place once a certain-sized hIAPP oligomer is formed. As we look toward higher order aggregates, questions arise about the formation of larger oligomers and whether these processes will proceed in a similar manner as dimer and 3-chain trimer formation, or perhaps a more complex process due to the increased number of monomers involved. Larger systems are being studied to further clarify these issues, and will be a target of future work.

## IV. METHODS

### A. Human Amylin Trimer

The hIAPP trimer system was designed based on the system used previously for hIAPP dimer simulations test by Guo, Fluitt, and de Pablo^19^. The amino acid sequence for hIAPP is KCNTATCATQRLANFLVHSSNNFGAILSSTNVGSNTY. Each C-termini is amidated, and Cys2 and Cys7 on each chain is linked by a disulfide bond. Protonation states were assigned on the basis of pKa values in water at a pH of 7.0; each hIAPP chain has a formal charge of +3, and chloride counterions are included to ensure charge neutrality. We use the AMBER ff99SB*-ILDN force field,^38–40^ which was chosen for its previously demonstrated ability to accurately capture behavior of amyloidogenic polypeptides and other intrinsically disordered proteins.^41,42^ The protein system was placed in a periodic cubic box with side length 15.0 nm with 110,008 TIP3P water molecules.^43^ Volume was kept constant, with coulombic forces calculated via the particle mesh Ewald algorithm^44,45^ and temperature held at 298 K using the Nosé-Hoover thermostat.^46^ A timestep of 2 fs was used, and hydrogen bond lengths were constrained to equilibrium values using the LINCS algorithm.^47^

### B. Finite Temperature String Method

We study hIAPP trimerization by employing the finite-temperature string method^48^, which calculates a transition pathway using a set of local points (“nodes”) connected in series by a smooth curve (“string”) in collective variable space. For the trimer system, we use two intuitive collective variables: (1) parallel *β*-sheet character 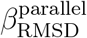, defined below in Equation 1; and (2) the radius of gyration *R_g_* of the three hIAPP chains, which provides a measure of spatial distance between each hIAPP monomer.

The parallel *β*-sheet character of a particular amino acid sequence between residues indices *u* and *v* is defined as:^49^

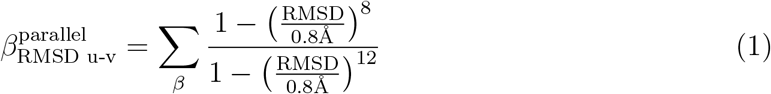

Equation 1 sums over every possible pair of three-residue segments bounded by residues *u* and *v* in each hIAPP monomer. “RMSD” refers to the root mean square deviation (in Å) of the positions of the N, C_α_, C*_β_*, C, and O backbone atoms of the residues in each pair of three-residue segments from those in an ideal parallel *β*-sheet. 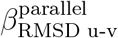 essentially measures the number of three-residue pairs that are arranged similarly to the configuration of an ideal parallel *β*-sheet.

Each string is discretized into 16 nodes, with each node’s location in collective variable space denoted by **z**_*α*_, where *α* is the node index along the string (*α* = 0,1, …, 15). The string nodes are used to generate a Voronoi tessellation, where each node is associated with a corresponding Voronoi cell, which consists of the region in CV space closer to its associated string node than any other node along the string. We assume Euclidian geometry for this collective variable space. At every iteration of the string method, each Voronoi cell is sampled such that there is no bias applied while the system is within the boundaries of the Voronoi cell; however, if the system departs from the Voronoi cell, a soft wall harmonic restraining potential is applied:

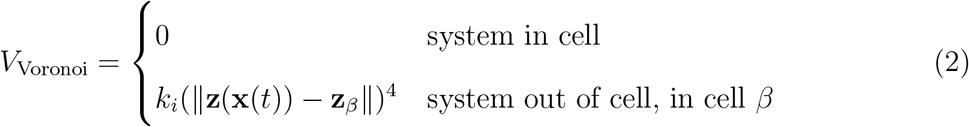

Each Voronoi cell is sampled for 100 ps per string method iteration, and the running average of each node’s explored location in collective variable space 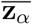 is tracked starting from the first string method iteration. The string is updated every nth iteration according to:

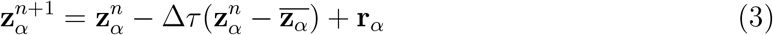

where we choose Δ*τ* to be 0.1, smoothing parameter *r_α_* to be 0 for nodes on each end of the string (*α* = 0 or 15), and for the interior nodes:

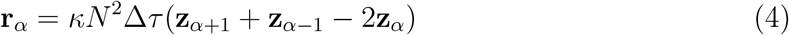

with smoothing parameter *κ* chosen to be 0.1 and the total number of nodes on the string *N* is 16. Following every string update, a cubic spline interpolation is drawn through the 16 nodes, and the nodes are then redistributed along the string in order to maintain equal arc-lengths between adjacent nodes. These steps are iterated until the string converges to a final pathway.

Upon convergence, the free energy is computed along the final string by calculating *π_α_*, the equilibrium probability of the system to be found in Voronoi cell *α*, which is then used to calculate the corresponding free energy *A_α_*^50^:

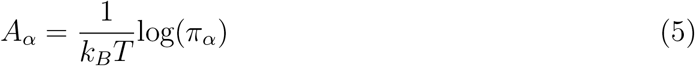

To improve resolution of the resulting free energy profile, we further discretize the original 16 node string to a total of *N* = 32 Voronoi cells. Each of the Voronoi cells is sampled using the same soft wall restraints described in Equation 2, for 50 ns for multiple runs. For each system sampling cell *α*, we collect *T_α_*, the total simulation time spent within cell *α*, as well as *N_αγ_*, the number of times the system escapes into a neighboring cell *γ*. The equilibrium probabilities *π_α_* are calculated with the following system of equations, where 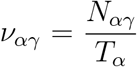 is the rate of escape from cell *α* into *γ*:

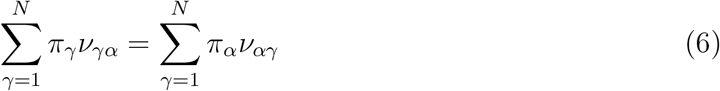

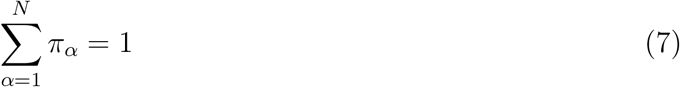

String method simulations were performed using the GROMACS 4.6.7 simulation package^51,52^, the PLUMED 2.1 plugin^53^, along with custom code to perform string method calculations.

## ACKNOWLEDGEMENTS

This work was completed using the computational resources of the University of Chicago Research Computing Center and the Laboratory Computing Resource Center at Argonne National Laboratory.

